# Environmental Microbial Community Signatures Associated with *Listeria* spp. Detection in German Meat Processing Facilities

**DOI:** 10.64898/2026.05.25.727608

**Authors:** Jérôme Braun, Nicole Wildi, Jasna Kovac, Claudia Guldimann

**Affiliations:** Competence Center for Food Safety, Chair of Food Safety and Analytics, LMU Munich, Oberschleißheim, Germany; Department of Food Science, The Pennsylvania State University, University Park, PA, USA; One Health Microbiome Center, Huck Institutes of the Life Sciences, The Pennsylvania State University, University Park, PA, USA

**Keywords:** *Listeria monocytogenes*, food processing environment, environmental microbiota, sampling, 16S rRNA amplicon sequencing, persistence, multilocus sequence typing

## Abstract

*Listeria monocytogenes* can persist in niches of meat processing environments despite routine cleaning and disinfection. Its persistence may depend not only on stress tolerance but also on interactions with resident microbial communities, which may promote or inhibit survival.

However, these ecological relationships remain poorly understood. We combined 16S rRNA V3/V4 amplicon sequencing, culture-based detection, and multilocus sequence typing (MLST) to characterize microbial communities in six German meat processing facilities over one year. We examined associations among community structure, sampling sites (drains and food-contact surfaces), and the occurrence of *Listeria* spp., including *L. monocytogenes*.

Microbial communities were dominated by core genera typical of food-processing environments, particularly *Pseudomonas* spp. and *Acinetobacter* spp., but differed significantly among facilities (PERMANOVA, p = 0.001; pairwise R² = 0.023–0.079), indicating facility-specific communities. Culture-based analyses detected *Listeria* spp. in 51 of 370 environmental samples (13.8%), mainly from drains (44/51, 86.3%). *L. monocytogenes* was detected in five of six facilities, with 19 of 21 isolates originating from drains (90.5%). MLST of 74 typeable *L. monocytogenes* isolates revealed high diversity, comprising 21 sequence types across 15 clonal complexes, with lineage II predominating (86.5%).

Overall microbial community composition was significantly associated with *Listeria* spp. and *L. monocytogenes* presence (PERMANOVA, p = 0.001; R² = 0.0137 and 0.0083). In drains, ASVs assigned to *Acinetobacter*, *Rhizorhapis*, and *Vagococcus* species showed positive associations with *Listeria* spp.-positive samples.

Together these findings suggest that drains are key ecological niches for *Listeria* spp. and that associated taxa may indicate drain communities linked to *Listeria* spp. recovery.

**IMPORTANCE:** *Listeria monocytogenes* is a major foodborne pathogen that can persist in meat processing environments despite routine cleaning and disinfection. Resident microbial communities may influence its survival, but longitudinal studies linking those communities with culture-based *Listeria* spp. detection remain limited. Here, we characterized microbial communities in six German meat processing facilities over 1 year using 16S rRNA gene amplicon sequencing and culture-based *Listeria* spp. detection, and MLST of *L. monocytogenes* isolates. We identified facility-specific microbial communities, identified floor drains as key niches for *Listeria* spp., and observed repeated recovery of different *L. monocytogenes* sequence types across facilities. In drains, ASVs assigned to the genera *Acinetobacter*, *Rhizorhapis*, and *Vagococcus* species were positively associated with culture-positive samples, identifying candidate taxa that may reflect microbial conditions associated with *Listeria* spp. recovery. These findings highlight the importance of considering not only whether *Listeria* spp. are detected, but also the resident microbial communities that may support their fitness.

## INTRODUCTION

*Listeria monocytogenes* is a foodborne pathogen of major public health concern due to its ability to persist in food-processing environments (FPEs). In humans, *L. monocytogenes* can cause listeriosis, a severe foodborne illness with high hospitalization and mortality rates and the highest fatality rate among the major zoonotic diseases reported in the EU in 2024 (European Food Safety Authority (EFSA), European Centre for Disease Prevention and Control (ECDC), 2025). In addition to its relevance for human health, *L. monocytogenes* is a major veterinary pathogen, underscoring the One-Health implications of this organism (Quereda et al., 2021). A key factor contributing to the persistence of *L. monocytogenes* in FPEs is its ability to grow at refrigeration temperatures and survive long-term in niches, particularly in biofilms (Fagerlund et al., 2017).

Other *Listeria* spp. are also frequently detected in food-processing environments and serve as index organisms for *L. monocytogenes* (Townsend et al., 2021).

However, physiological resilience alone does not fully explain persistence. Increasing evidence indicates that ecological interactions within resident microbial communities play a critical role in microbial fitness. Biofilms formed by dominant colonizers, such as *Pseudomonas* spp., can enhance the resilience of *L. monocytogenes* against disinfectants (Rolon et al., 2024; Voloshchuk et al., 2025), while other community members may modulate survival through nutrient competition, inhibitory metabolites, or changes in biofilm structure and local microenvironmental conditions (Fagerlund et al., 2021).

Stable, facility-specific microbial communities have been described across diverse FPEs, including breweries, fruit processing facilities, and meat processing facilities (Belk et al., 2022; Bokulich et al., 2015; Rolon et al., 2023). These resident microbiota form persistent assemblages shaped by the environment, raw materials, processing steps, and cleaning regimens (Barcenilla et al., 2024). Until recently, the microbial communities of meat processing environments were comparatively underexplored, and substantial knowledge gaps remained regarding their composition, stability, and ecological drivers (Belk et al., 2022; Burnett et al., 2025). Recent metagenomic and longitudinal studies have started to address these gaps, revealing that meat processing environments harbor diverse, facility-specific, and persistent microbial communities, including previously undescribed taxa and biofilm-forming strains (Barcenilla et al., 2024).

Understanding these microbial ecosystems is essential for identifying how individual community members may facilitate or inhibit the persistence of *L. monocytogenes* (Fagerlund et al., 2017).

This knowledge may reveal antagonistic interactions or microbiota-derived inhibitory compounds that could be exploited for preventive purposes and for the development of targeted hygiene strategies (Belk et al., 2022; Zwirzitz et al., 2021).

In this study, we characterized the microbiota of six German meat processing facilities over a one-year period and investigated the composition of the microbial communities and their associations with the occurrence of *Listeria* spp. and *L. monocytogenes.* We combined longitudinal 16S rRNA gene sequencing with culture-based detection of *Listeria* spp. and *L. monocytogenes* to determine whether specific microbial community signatures are associated with *Listeria* spp. occurrence and persistence in meat processing environments. Multilocus sequence typing (MLST) was used to subtype *L. monocytogenes* isolates and identify patterns of repeated recovery indicative of persistence. Beyond characterizing facility-specific microbial communities, we identified taxa whose abundance or prevalence was positively or negatively associated with culture-based *Listeria* spp. detection. These taxa provide an ecological context for *Listeria* spp. occurrence and represent candidates for future experimental interaction studies. Together, these analyses link microbiome composition with *Listeria* spp. occurrence and environmental reservoirs in meat processing environments.

## MATERIALS AND METHODS

### Sampling and sample processing

Samples were collected from six German meat processing facilities and covered all major production areas, including receiving, cold storage, raw material handling, and finished product processing. Per area, at least one floor drain and one food-contact surface (FCS) were sampled, resulting in 12 sites per facility (14 in one larger facility). The facilities processed either beef (F5), pork (F2, F3, F4, F6), or both (F1). Each facility was sampled five times at ∼10-week intervals. Sampling was conducted on Mondays before production started to allow for the recovery of the microbiota during the weekend after routine cleaning and disinfection on Fridays. At each site, two adjacent but non-identical areas were sampled, yielding 740 samples (370 paired swabs): one for microbiota profiling (16S rRNA V3/V4 sequencing) and one for culture-based detection of *Listeria* spp., including *L. monocytogenes*, according to ISO 11290-1:2017, with species-level identification by MALDI-TOF MS. For both FCS and drains, ∼225 cm² were sampled using sponge-sticks with neutralization buffer (Neogen Corporation, Lansing, MI, USA). If present, standing water was additionally collected from drains. All samples were cooled during transport to the lab and processed within 12 h.

For the samples intended for sequencing, sponges were aseptically transferred into sterile VWR Blender Bags (Filter Bag 129-0734; Avantor, Radnor, PA, USA) containing an integrated filter membrane, homogenized in 50 mL buffered peptone water (Merck KGaA, Darmstadt, Germany) for 30 s at 230 rpm (Stomacher 400 Circulator, Seward Ltd, Worthing, West Sussex, UK), and the filtered homogenate was split into two 25-mL aliquots. Both aliquots were centrifuged (16,100 × g, 35 min, room temperature) to obtain bacterial pellets. One pellet was used for 16S rRNA V3/V4 gene sequencing, while the other was resuspended in 1 mL Brain Heart Infusion broth supplemented with 15% glycerol and stored at -80 °C.

### DNA Extraction

For 16S rRNA V3/V4 gene sequencing, bacterial DNA was extracted from the pellet using the DNeasy Blood & Tissue Kit (Qiagen, Hilden, Germany) following the manufacturer’s protocol, including the pre-treatment step for Gram-positive bacteria. With each DNA extraction batch, one positive extraction control (mock microbial communities from ZymoBIOMICS Microbial Community Standard D6300; Zymo Research Europe GmbH, Freiburg, Germany) and one negative extraction control were included to assess extraction performance and monitor potential contamination. Negative extraction controls consisted of clean sponges that were processed in bags in the same way as the samples.

Samples were eluted in 50 µL of preheated Buffer ATE (65 °C). DNA concentration was subsequently quantified using the Quantus Fluorometer with QuantiFluor dsDNA Dye (Promega, Madison, WI, USA). DNA was stored at -20 °C until sequencing. Library preparation for V3/V4 16S rRNA gene sequencing was done using barcoded 341F (forward) and 806R (reverse) primers (Klindworth et al., 2013) by Novogene GmbH (Novogene, Munich, Germany). The PCR reactions were carried out with 15 μL of Phusion High-Fidelity PCR Master Mix; 0.2 μM of forward and reverse primers, and 10 ng template DNA. Thermal cycling consisted of initial denaturation at 98 ℃ for 1 min, followed by 30 cycles of denaturation at 98 ℃ for 10 s, annealing at 50℃ for 30 s, and elongation at 72℃ for 30 s, followed by a final 5-min hold at 72℃. Library preparation was done by Novogene GmbH (Novogene, Munich, Germany) using the NEBNext® Ultra™ II DNA Library Prep Kit (New England Biolabs, Ipswich, MA, USA) and sequencing was performed on an Illumina NovaSeq 6000 (PE 250 bp reads, 50k reads/sample).

### Sequencing data analyses

Sequencing data analyses were carried out in RStudio, R v. 4.5.0 (R Core Team, 2025) unless otherwise noted, using the DADA2 package v. 1.36.0 (Callahan et al., 2016). Demultiplexed paired-end reads were analyzed as outlined in the DADA2 analysis.R script available on GitHub (see Data Availability). Briefly, sequences were inspected for quality and used to learn error rates. Amplicon sequence variants (ASVs) were inferred using DADA2. Forward and reverse reads were merged, sequences of non-target length were removed, and chimeras were identified and removed using the consensus method. Taxonomic assignment was performed using the SILVA v138.1 reference database (Quast et al., 2013). Negative extraction controls were used for contaminant identification using prevalence-based contaminant identification with decontam package v. 1.30.0, as outlined in the decontam.R script available on GitHub (Davis et al., 2018) in R version 4.5.1. Identified contaminant ASVs were removed prior to statistical analyses.

### Statistical analysis of microbial community data

Statistical analyses were conducted as outlined in the following sections and in the statistical_analysis.R script provided on GitHub.

### Beta diversity analysis

To account for the compositional nature of data, zero counts in the ASV table were imputed using the Count Zero Multiplicative (CZM) approach with zCompositions package v. 1.5.0.5 (Palarea-Albaladejo and Martín-Fernández, 2015). Centered log-ratio (CLR) transformation was then applied to the filtered ASV table prior to principal component analysis (PCA) and plotting of PCA biplots to visualize clustering of samples. To statistically assess differences in beta diversity (i.e., microbiota composition), Aitchison distances were computed, and permutational multivariate analysis of variance (PERMANOVA) was conducted using the pairwiseAdonis package v. 0.4.1 (Martinez Arbizu, 2020). Models tested main effects and interactions among facility, sampling area, *Listeria* spp. and *Listeria monocytogenes* presence, with 999 permutations. p-values were Bonferroni-adjusted.

### Differential abundance analysis

MaAsLin3 v. 1.1.2 was used to identify ASVs associated with *Listeria* spp. or *L. monocytogenes* (Nickols et al., 2026). Analysis was applied to all samples, as well as to a subset of samples from drains, due to more frequent recovery of *Listeria* spp. from drain samples. The analyses were conducted using an untransformed ASV table and total-sum scaling (TSS). Associations with a false discovery rate-adjusted significance threshold of 0.1 were considered for interpretation.

### Core microbiota analysis

Core microbiota were defined at the genus level using untransformed relative abundance data. For each facility, genera present in at least 90% of samples at a minimum relative abundance of 0.0001 were identified as core genera, and their mean relative abundance and standard deviation were calculated. Overall core genera were also identified, using the same criteria, for all samples collected in this study. Further, the ten most abundant genera per facility were plotted using ggplot2 v. 4.0.0 (Wickham, 2016) to visualize the most abundant genera in each facility.

### Alpha diversity analysis

Alpha diversity was assessed using the Shannon index calculated from the contaminant-filtered ASV count table after rarefaction to the minimum sequencing depth of 23,689 reads per sample. Rarefaction was performed using the rrarefy function in the vegan package v. 2.7.3 to account for differences in sequencing depth among samples, and Shannon diversity was calculated from the rarefied ASV count table using the diversity function in vegan. For longitudinal analyses, sampling sites were defined as repeatedly sampled locations identified by the sampling site code within each facility. The R code used for alpha diversity and longitudinal community-change analyses is provided in longitudinal_analysis.R in the GitHub repository listed under Data Availability.

### Longitudinal community change

Temporal changes were quantified as the mean Aitchison distances in microbial community composition between samples collected from the same sampling unit at consecutive sampling events. Sampling units were defined by the sampling site code. Facility-stratified temporal trends in Shannon diversity and longitudinal community change were analyzed using linear mixed-effects models fitted with lme4 v. 2.0.1 and lmerTest v. 3.2.1. Models included sampling date as a numeric fixed effect and sampling site, or sampling unit for longitudinal community change, as a random intercept. Model outputs were extracted using broom.mixed v. 0.2.9.7, and p-values from facility-stratified models were adjusted using the false discovery rate method.

### Sequencing and culture concordance

To assess the association between sequencing-based *Listeria* spp. signal and culture-based *Listeria* spp. detection, a logistic regression model was fitted using culture positivity as the binary response variable and *Listeria* species ASV abundance as the predictor.

### *Listeria* spp. Enrichment

Samples collected for culture-based detection of *Listeria* spp. and *L. monocytogenes* were processed according to ISO 11290-1:2017 to detect viable *Listeria* spp., as sequencing-based methods cannot differentiate viable from dead bacteria. Briefly, sample homogenates were subjected to primary enrichment in half-Fraser broth, followed by secondary enrichment in Fraser broth. After incubation, aliquots from the enrichment cultures were streaked onto Chromocult *Listeria* Agar according to Ottaviani and Agosti and GranuCult Oxford Agar, each supplemented with the respective *Listeria*-selective supplement according to the manufacturer’s instructions (all Merck KGaA, Darmstadt, Germany). Selective agar plates were evaluated after 48 h of incubation. For each positive sample, one colony showing morphology typical of *Listeria* spp. on the ALOA plate was subcultured on plate count plates for 24 h and subjected to MALDI-TOF MS identification to confirm the species. Colonies were prepared using a formic acid extraction protocol: bacterial material from a single colony was suspended in 300 µL deionized water, ethanol was added, and the suspension was centrifuged. The dried pellet was extracted with 70% formic acid and acetonitrile, and 1 µL of the supernatant was spotted onto a MALDI target plate, dried, overlaid with 1 µL HCCA matrix solution, and analyzed using the MALDI Biotyper Sirius system with the BDAL database version 2023 (Bruker Daltonics, Bremen, Germany). When colonies with different morphologies suggestive of *Listeria* spp. were observed from the same enriched sample, representative colonies of each morphology were subcultured separately and identified by MALDI-TOF MS. Thus, multiple *Listeria* species could be obtained from a single culture-positive environmental sample.

### MLST of Listeria monocytogenes Isolates

MLST was used to subtype *L. monocytogenes* isolates, assign sequence types and clonal complexes, and assess patterns of persistence within and across facilities. It was performed according to the standardized scheme (Ragon et al., 2008; Salcedo et al., 2003), targeting seven housekeeping genes of *L. monocytogenes abcZ, bglA, cat, dapE, dat, ldh, and lhkA*.

Isolates originated from environmental samples collected during the present study and from 55 additional isolates provided by the participating facilities as part of their routine environmental monitoring conducted between 2020 and 2025. Twelve of the latter were collected outside the sampling period of the present study (2024-2025).

Genomic DNA was extracted by heat lysis based on a previously described protocol (Dashti et al., 2009), with minor modifications. For each isolate, a single colony was inoculated into 5 mL Brain Heart Infusion broth and incubated for 24 h. The entire overnight culture was centrifuged to obtain a cell pellet. Cell pellets were resuspended in 250 μL molecular-grade water, incubated at 99°C for 10 min, and centrifuged at 5000 × g for 5 min. Then, 200 μL of the supernatant was transferred to a clean microcentrifuge tube and used as a DNA template in PCR amplification.

PCR amplification was performed using SensiFAST SYBR No-ROX Master Mix (Bioline, London, UK) on a Bio-Rad CFX96 real-time PCR system. Each 20 µL reaction contained 10 µL 2× SensiFAST SYBR No-ROX Master Mix, forward and reverse primers at a final concentration of 0.4 µM each, 7.4 µL molecular-grade water, and 1 µL template DNA diluted 1:1000 for each sample before amplification. The same primer concentration was used for all MLST targets.

Cycling conditions followed the Institut Pasteur MLST recommendations: 95 °C for 5 min; 40 cycles of 95 °C for 10 s, 45 or 52 °C for 10 s, and 72 °C for 20 s; followed by 72 °C for 2 min. An annealing temperature of 45 °C was used for *bglA, cat*, and *ldh,* and 52 °C for all other targets.

Sanger sequencing was performed by Eurofins Genomics (Ebersberg, Germany). The loci *abcZ*, *cat*, *dat*, and *ldh* were sequenced in a single direction using the forward primer only, while *bglA*, *dapE*, and *lhkA* were sequenced bidirectionally using both forward and reverse primers.

Sequencing chromatograms were imported into Geneious Prime version 2025.0.3 for quality assessment. Chromatograms were inspected for signal intensity, base-calling accuracy, and the presence of mixed or ambiguous peaks. For bidirectionally sequenced loci, forward and reverse reads were assembled and used to generate consensus sequences. High-quality single-direction sequences and bidirectional consensus sequences were exported as FASTA files and uploaded to the Institut Pasteur BIGSdb-Lm database (Institut Pasteur, n.d.) for allele designation and sequence type (ST) assignment according to the canonical seven-locus MLST scheme.

Persistence was defined as the repeated isolation of *Listeria monocytogenes* isolates belonging to the same ST within the same facility at sampling times >6 months apart. This conservative time-based criterion was adopted because no universally accepted definition of *Listeria* persistence in food-processing environments currently exists (Belias et al., 2022). Repeated recovery of the same ST at intervals <6 months was described as recurrent detection.

## RESULTS

### Controls indicated reproducible sequencing performance and low contamination

Mock community samples showed consistent results across runs, with all expected standard taxa recovered (Fig. S1). The observed relative abundances did not fully match the theoretical composition due to lower extraction efficiency for *Staphylococcus aureus,* suggesting an extraction bias against this species and potentially other Gram-positive taxa. Negative controls produced zero reads in 11 out of 16 cases, rare low-read contaminants were identified in 5 samples (mean: 1309.4 reads) and used as input for the *decontam* pipeline (Davis et al., 2018).

### Distinct facility microbial communities shared core genera of *Pseudomonas* and *Acinetobacter*

Among the 370 samples submitted for 16S rRNA gene sequencing, 366 generated sufficient sequencing reads for downstream analysis, while four samples yielded no reads and were therefore excluded from subsequent microbiota analyses.

Microbial community composition differed significantly among facilities (Fig. 1). Pairwise PERMANOVA comparisons showed that all facilities differed significantly from each other (p = 0.001), but the proportion of variation explained by facility differed among pairwise comparisons. The largest effect sizes were observed for comparisons involving F6, particularly F1 vs. F6 (R² = 0.079), F4 vs. F6 (R² = 0.076), F5 vs. F6 (R² = 0.062), and F6 vs. F2 (R² = 0.062). In contrast, the smallest effect size was observed for F3 vs. F2 (R² = 0.023).

**FIG 1.**
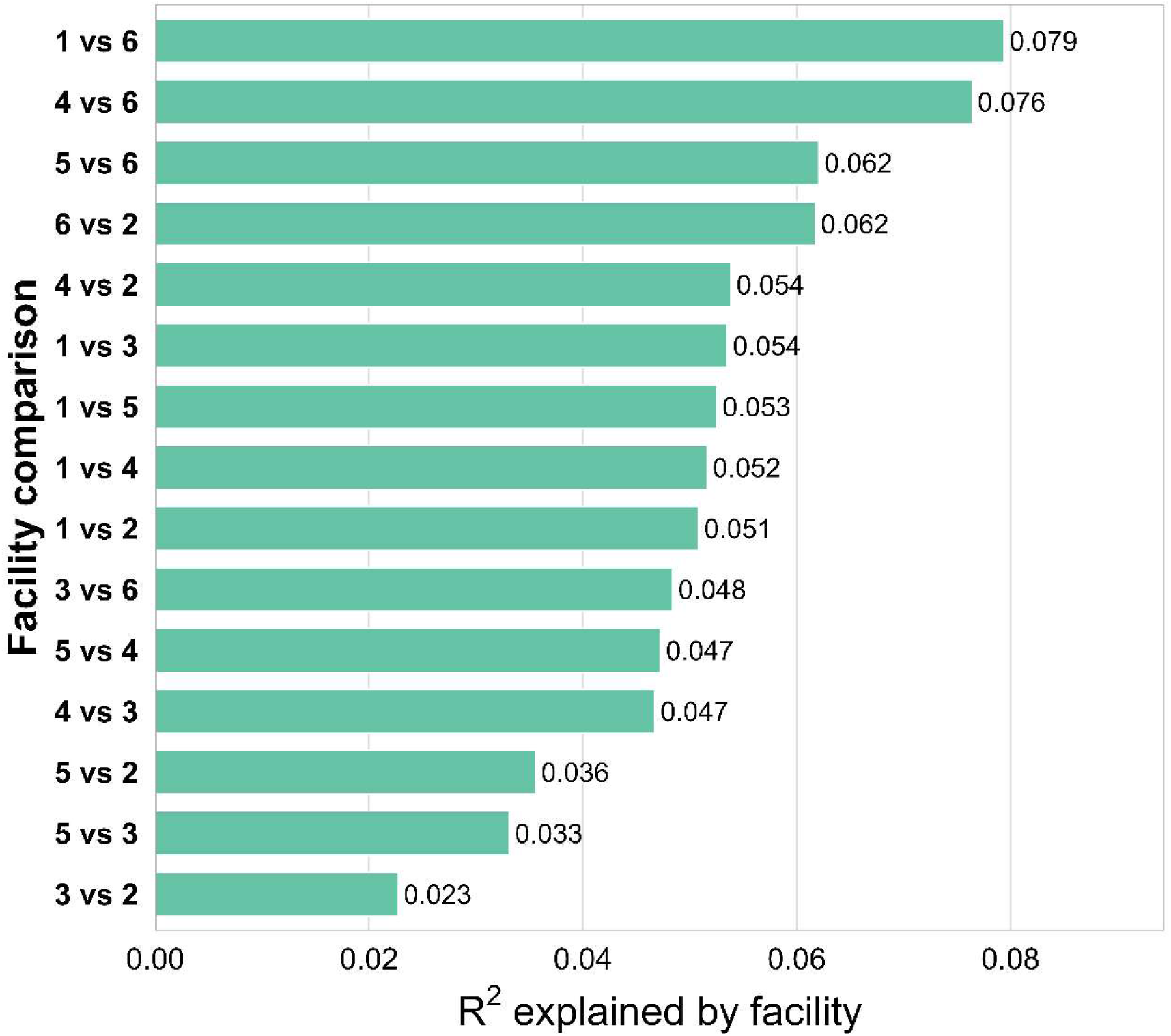
Pairwise PERMANOVA effect sizes (R²) for microbial community composition between meat processing facilities. Bars show the proportion of variance explained by facility (R²) for each pairwise comparison between the six meat processing facilities. All pairwise comparisons were significant (PERMANOVA, p = 0.001), indicating significant differences in microbial community composition among facilities.

Genus-level relative abundance profiles varied across facilities and sampling times (Fig. 2). *Pseudomonas* spp. and *Acinetobacter* spp. were repeatedly detected at high relative abundance in all six facilities, but their relative contributions varied across facilities. *Pseudomonas* spp. was prominently present in several facilities, whereas *Acinetobacter* spp. was particularly abundant in F3 and F5. Consistent with their broad distribution, *Pseudomonas* and *Acinetobacter* were the only genera identified as core genera across all samples, occurring in more than 90% of samples at a relative abundance of at least 0.0001. Their mean relative abundances were 0.17 ± 0.24 and 0.16 ± 0.22, respectively.

**FIG 2.**
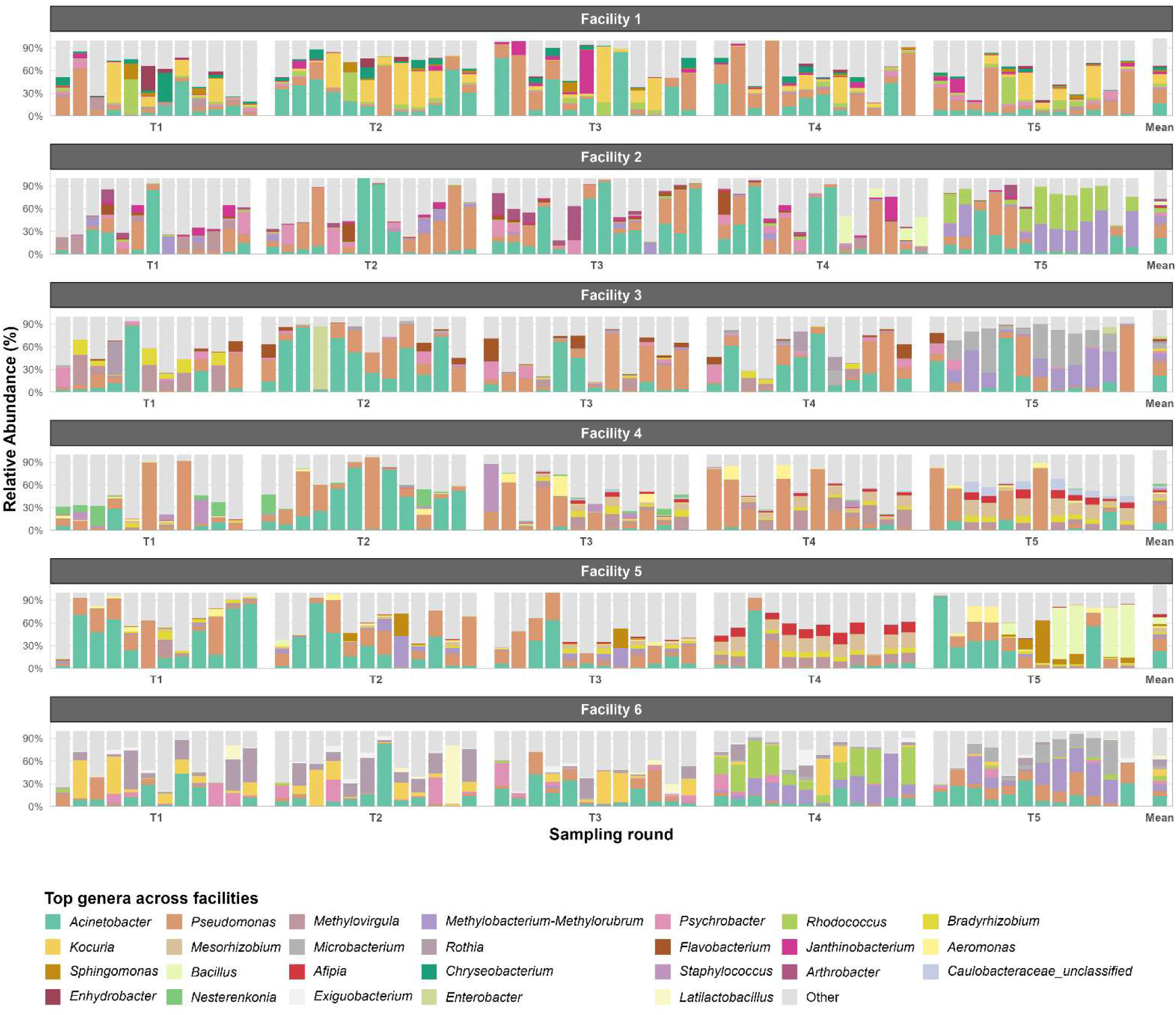
Relative abundance of genera in six meat processing facilities over time. Stacked bar plots show the relative abundance of bacterial genera in individual samples collected from six meat processing facilities across five sampling times (T1–T5). Each bar represents one sample. For each facility, the ten genera with the highest mean relative abundance across all samples from that facility are shown individually; all remaining genera are grouped as “Other”. The bar labeled “Mean” (far right) represents the mean relative abundance of these genera across all samples from the respective facility.

Core genera were defined as genera present in at least 90% of samples from a given facility at a minimum relative abundance of 0.0001. Facility-specific core genera differed among facilities (Fig. 4) and overlapped with several of the dominant genera shown in the facility-level mean profiles in Fig. 3. In F2, *Pseudomonas* spp. and *Acinetobacter* spp. were the only core genera, whereas the remaining facilities contained additional facility-specific core genera. These included *Chryseobacterium*, *Janthinobacterium*, *Kocuria*, and *Rhodococcus* species in F1; *Microbacterium* spp. in F3; *Aeromonas* spp. in F4 and F5; *Chryseobacterium*, *Flavobacterium*, and *Janthinobacterium* species in F5; and *Enhydrobacter*, *Exiguobacterium*, *Kocuria*, *Microbacterium*, *Paracoccus*, *Psychrobacter*, *Rhodococcus*, and *Rothia* species in F6. Several facility-specific core genera, including *Rhodococcus*, *Psychrobacter*, *Flavobacterium*, and *Chryseobacterium* species, were also among the abundant genera shown in Fig. 3.

**FIG 3.**
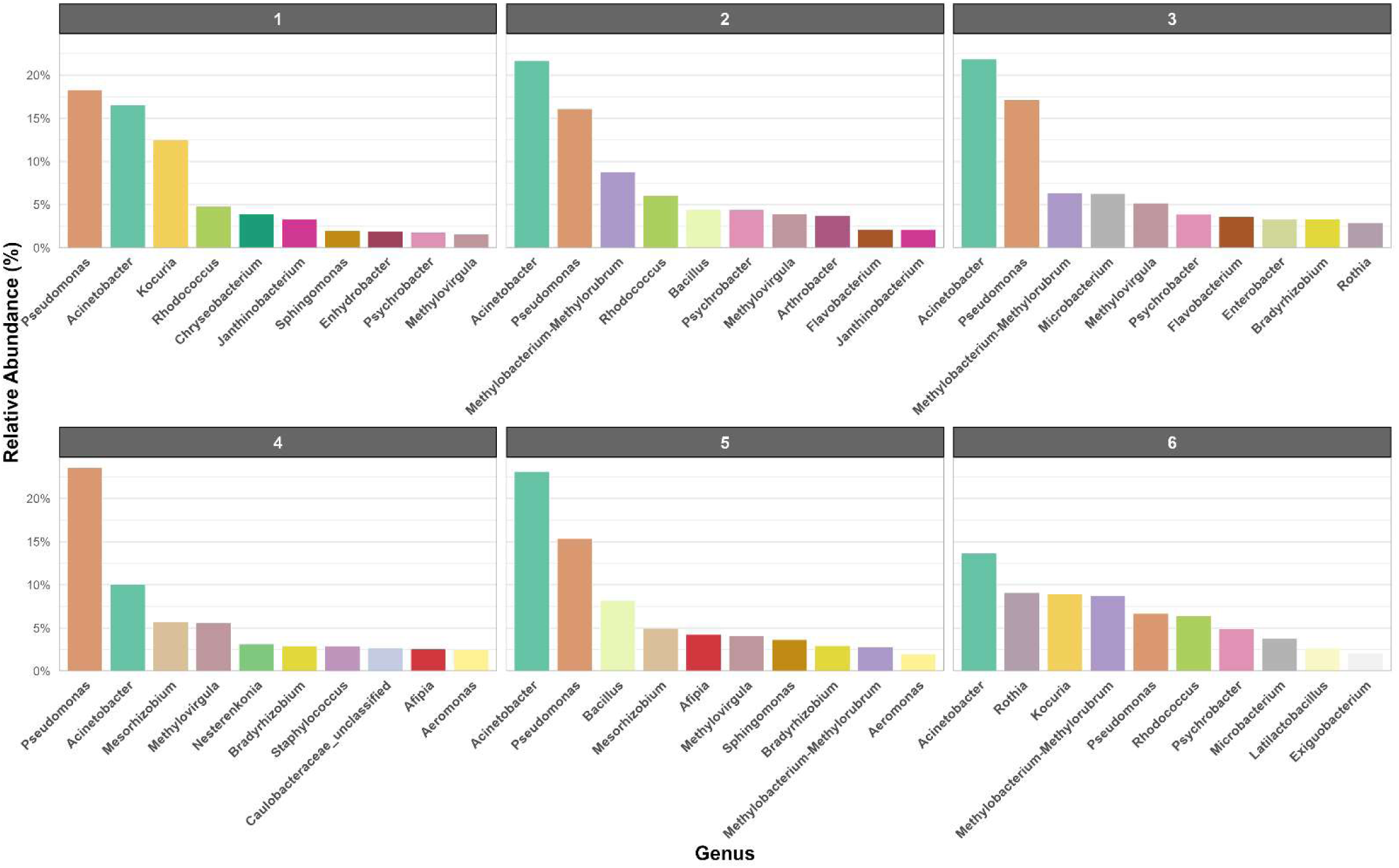
Ten genera with the highest relative abundance in each facility. This figure provides an expanded view of the rightmost “mean” bars shown in Fig. 2. For each facility, the mean relative abundance of the ten genera with the highest mean relative abundance across all samples is displayed.

**FIG 4.**
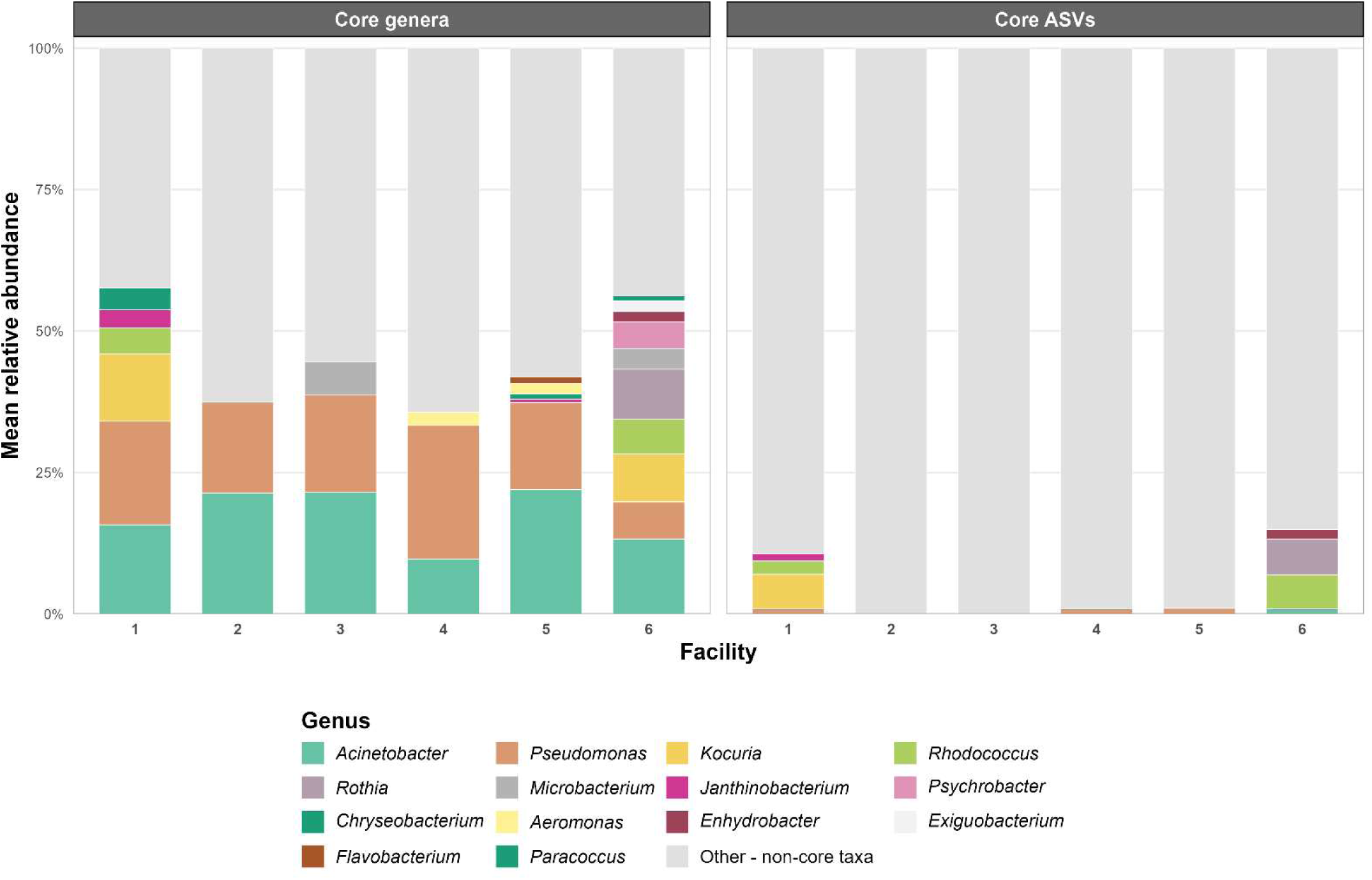
Facility-specific composition of core taxa (≥90% prevalence, ≥0.0001 relative abundance). Bars show the mean relative abundance of facility-specific core taxa across samples from each facility, with all remaining taxa grouped as “Other – non-core taxa”. The left panel shows core genera, whereas the right panel shows core ASVs aggregated to genus level for visualization. Genus colors were kept consistent across panels to facilitate comparison.

At the ASV level, fewer core taxa were identified than at the genus level, and core ASVs differed among facilities (Fig. 4). Using the same prevalence and relative abundance criteria as above, no core ASVs were identified in F2 or F3, indicating lower ASV-level core stability in these facilities. F4 and F5 each contained one core ASV (ASV42 and ASV16, respectively), both assigned to *Pseudomonas* spp., which was among the ten most abundant genera in these facilities. In F1 and F6, several core ASVs were identified. In F1, core ASVs were assigned to *Rhodococcus* ASV1, *Kocuria* ASV7, *Janthinobacterium* ASV26, and *Pseudomonas* ASV64, whereas in F6, core ASVs were assigned to *Rhodococcus* ASV1, *Rothia* ASV11, *Acinetobacter* ASV14, and *Enhydrobacter* ASV25. These genera were also represented in the facility-level dominant genus profiles shown in Fig. 3.

### Temporal dynamics of microbial communities

Alpha diversity, as measured by the Shannon index, showed temporal variation across the study period (Fig. 5). Shannon diversity varied significantly across sampling events (LRT, q = 7.74 × 10⁻¹³), whereas production area was not significantly associated with Shannon diversity (q = 0.419). However, the interaction between production area and sampling event was significant (q = 0.025), indicating that temporal patterns in Shannon diversity differed among production areas. In addition, the categorical sampling-event model provided a significantly better fit than a model assuming a linear time effect (q = 2.00 × 10⁻¹⁰).

**FIG 5.**
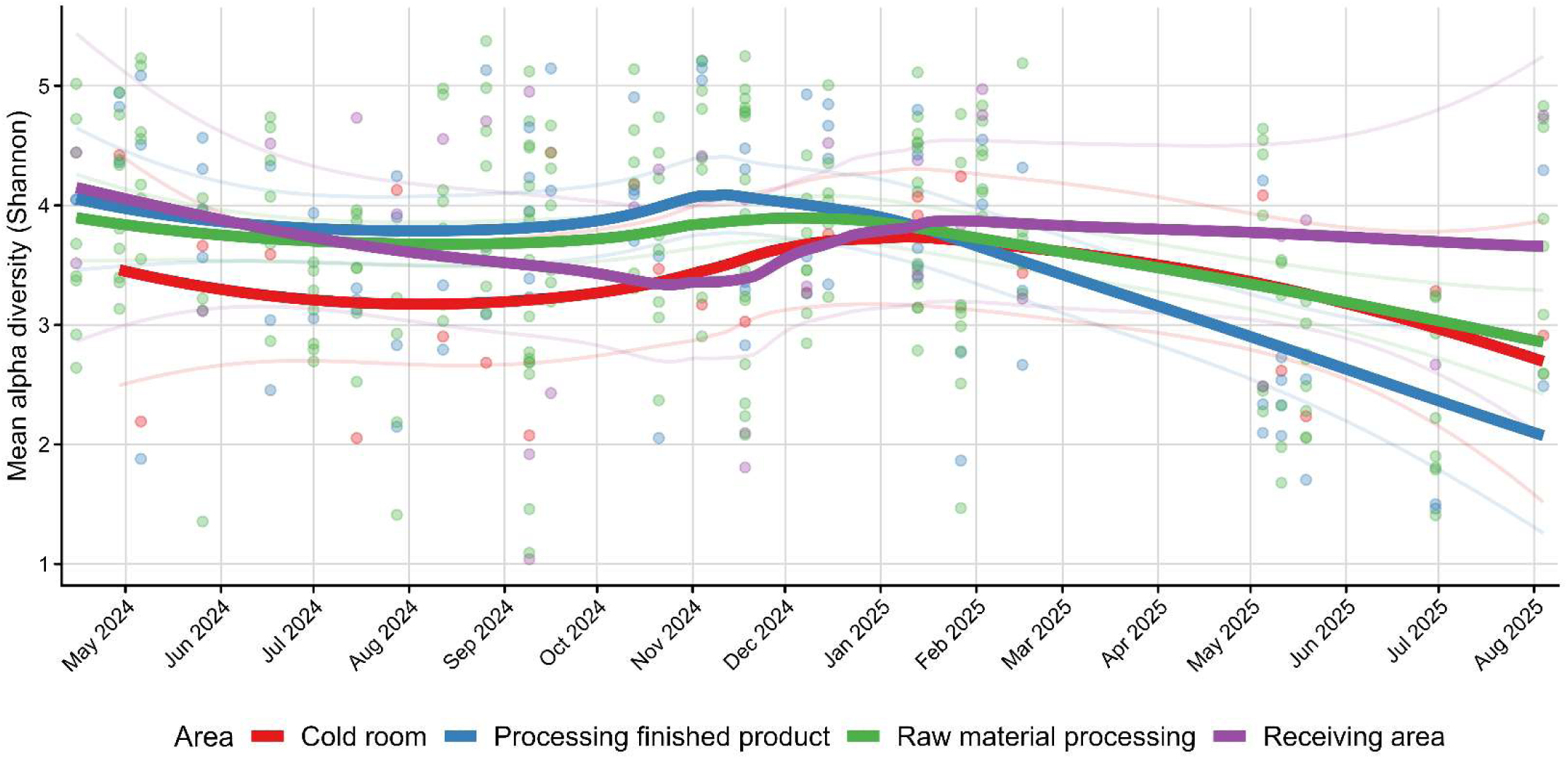
Temporal dynamics of alpha diversity across production areas. Shannon diversity is shown for samples collected across the study period. Points represent individual samples and are colored by production area. Thick colored lines indicate area-level mean curves, and thin colored lines show the upper and lower limits of the 95% confidence intervals. Sampling event was modeled as a categorical predictor to assess temporal variation without assuming a linear trend.

As a measure for beta diversity, longitudinal community change, quantified as the Aitchison distance from the previous sampling event was reported. It varied significantly across sampling events (LRT, q = 8.29 × 10⁻⁸; Fig. 6). In contrast to Shannon diversity, production area was significantly associated with longitudinal community change after accounting for sampling event (q = 0.022), indicating that the magnitude of community turnover differed among production areas. The interaction between production area and sampling event was also significant (q = 0.0099), indicating that temporal patterns in community change differed among production areas. Consistent with the non-monotonic mean curves, the categorical sampling-event model provided a significantly better fit than a model assuming a linear time effect (q = 8.29 × 10⁻⁸).

**FIG 6.**
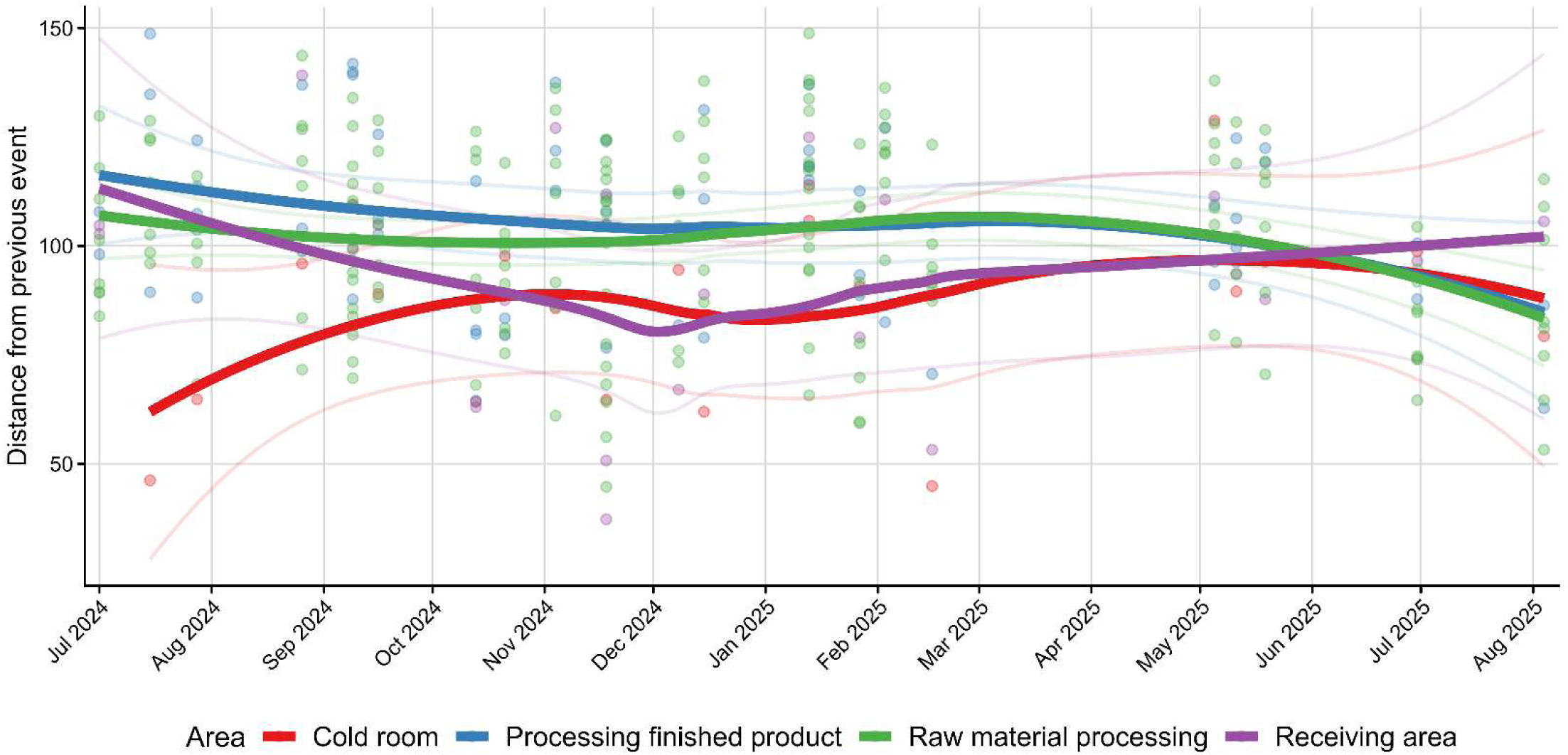
Longitudinal community change across production areas. Longitudinal community change was quantified as the Aitchison distance between samples collected from the same sampling unit at consecutive sampling events. Points represent individual sampling units and are colored by production area. Thick colored lines indicate area-level mean curves, and thin colored lines show the upper and lower limits of the 95% confidence intervals. Sampling event was modeled as a categorical predictor to assess temporal variation without assuming a linear trend.

Facility-stratified mixed-effects models were refitted, with the sampling event modeled as a categorical predictor, to assess temporal variation without assuming a linear trend. Shannon diversity varied significantly across sampling events in five of six facilities after FDR correction, namely facilities F1, F2, F3, F5, and F6 (q = 1.0 × 10⁻⁵ to 0.0367; Fig. S2), whereas no significant temporal variation was detected in facility 4 (q = 0.181).

Longitudinal community change, measured as the Aitchison distance from the previous sampling event, varied significantly over time in facilities F1, F2, F4, and F6 (q = 1.5 × 10⁻⁴ to 0.0262; Fig. S3), while facilities F3 and F5 did not show significant temporal variation after FDR correction. These results indicate that temporal microbiome dynamics were facility-specific and not consistently monotonic across the study period.

### *Listeria* spp. and *L. monocytogenes* were significantly more likely to be detected in drains than on food-contact surfaces

Of 370 environmental samples, 51 (13.8%) were positive for *Listeria* spp. Because multiple *Listeria* spp. were recovered from some positive samples, these 51 positive samples yielded 58 isolates representing six species (Table 1). *L. monocytogenes* was detected in five facilities, with the highest numbers in F2 and F3. Most *L. monocytogenes* isolates from the present survey were recovered from drains (19/21), whereas only two were recovered from FCS. Overall, *Listeria* spp.-positive samples were significantly more frequent in drains than on FCS (44/180 vs. 7/190; Fisher’s exact test, p = 2.55 × 10⁻⁹; OR = 8.46). Co-occurrence of multiple *Listeria* spp. was observed in 7 out of 51 positive samples (13.7%), most commonly involving *L. monocytogenes* and *L. innocua* (5/7).

**Table 1.** Distribution of *Listeria* species across the six processing facilities. Numbers represent positive isolates per facility and species. Values in brackets indicate food-contact-surface-positive samples. All other positive samples were obtained from drains.

| Species | Facility |  |  |  |  |  | Total |
| --- | --- | --- | --- | --- | --- | --- | --- |
|  | F1 | F2 | F3 | F4 | F5 | F6 |  |
| *L. monocytogenes* | 1 | 6 (2) | 6 | 0 | 4 | 4 | 21 (2) |
| *L. innocua* | 0 | 4 (1) | 8 | 1 | 1 | 7 (2) | 21 (3) |
| *L. welshimeri* | 4 (1) | 2 (1) | 3 | 1 | 0 | 0 | 10 (2) |
| *L. grayi* | 0 | 0 | 0 | 2 | 1 | 0 | 3 |
| *L. seeligeri* | 0 | 0 | 0 | 0 | 0 | 2 | 2 |
| *L. newyorkensis* | 0 | 0 | 0 | 0 | 0 | 1 | 1 |
| **Total isolates** | **5 (1)** | **12 (4)** | **17 (0)** | **4 (0)** | **6 (0)** | **14 (2)** | **58 (7)** |

### Microbial community composition differed significantly between *Listeria-*spp. positive and *Listeria-*spp. negative samples, with specific ASVs associated with *Listeria* spp.-positive samples

Logistic regression revealed no significant association between *Listeria* ASV abundance and culture-based detection of *Listeria* spp. (estimate = -0.0012 ± 0.00086, p = 0.162), indicating limited concordance between sequencing-based *Listeria* spp. signal and culture positivity.

At the community level, the presence of *Listeria* spp. and *L. monocytogenes* were associated with minor but significant shifts in microbiota composition (PERMANOVA: R² = 0.0137 and 0.0083, respectively; both p = 0.001). Since *Listeria* spp.-positive samples were predominantly recovered from non-food-contact surfaces (i.e., drains), we conducted further differential abundance analyses on a subset containing only the drain samples (Fig. 7). In drain samples, three ASVs were significantly associated with culture-based *Listeria* spp. status at an FDR threshold of 0.1.

**FIG 7.**
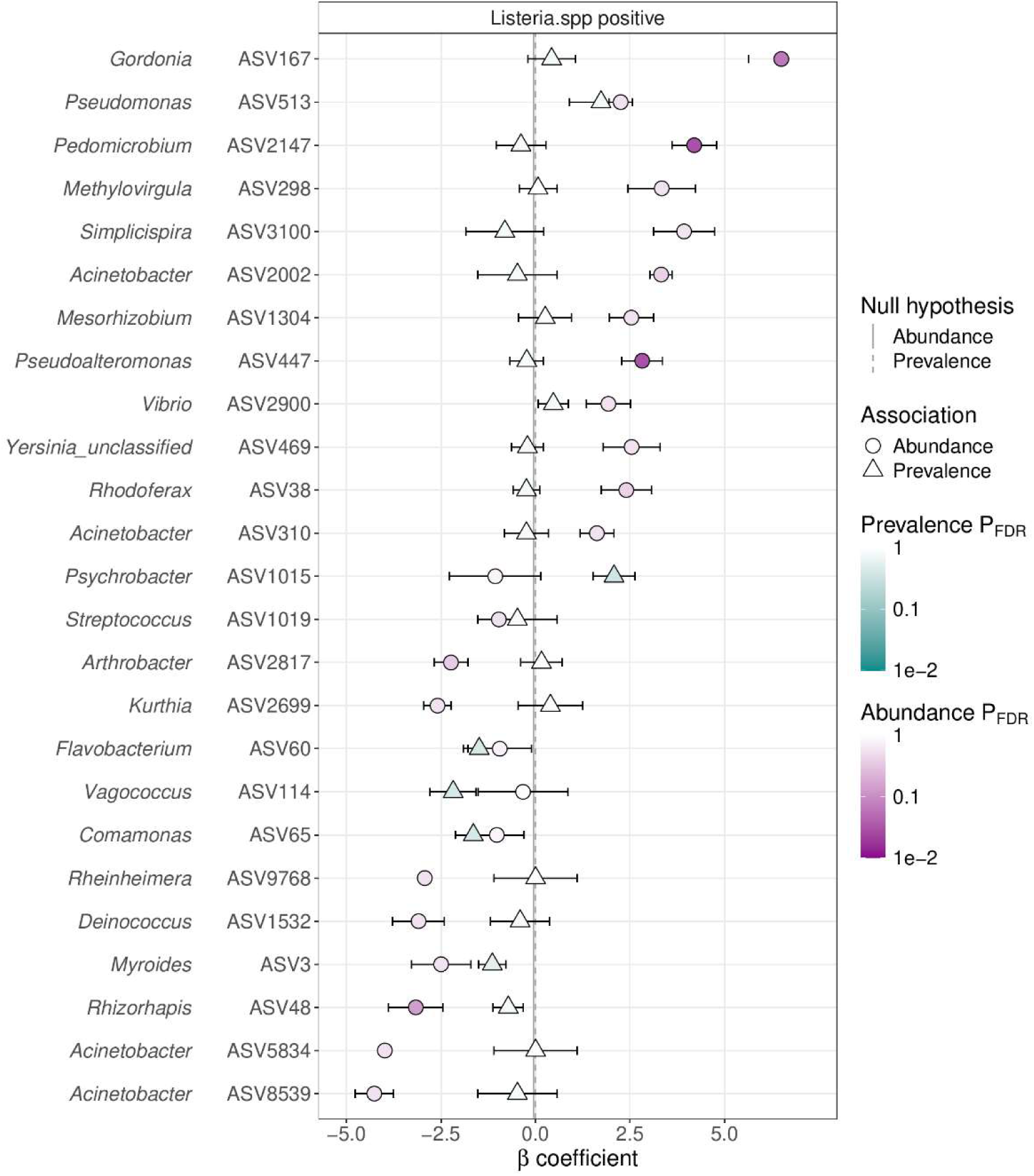
ASVs associated with *Listeria* spp. in drain samples. Differential abundance of ASVs (circles) and log-odds of ASV presence (triangles) in *Listeria* spp.-positive samples are shown. Positive β coefficients indicate positive associations with *Listeria* spp.-positive drain samples, whereas negative β coefficients indicate negative associations. Associations were considered significant at an FDR-adjusted threshold of q < 0.1.

All three were detected in the abundance model and were positively associated with *Listeria* spp.-positive samples: *Acinetobacter* ASV167 (β = 6.50, q = 0.064), *Rhizorhapis* ASV2147 (β = 4.19, q = 0.029), and *Vagococcus* ASV447 (β = 2.82, q = 0.029). No ASVs were significantly associated in the prevalence model, and no ASVs showed significant negative associations after FDR correction. In contrast, no ASVs were significantly associated with culture-based *L. monocytogenes* status in drain samples in either the abundance or prevalence model after FDR correction.

### MLST revealed high genetic diversity and persistent sequence types

For MLST analysis, a total of 76 *L. monocytogenes* isolates were included, comprising 21 isolates recovered from environmental samples collected during the present study and 55 additional isolates provided by the participating facilities from their routine monitoring programs. They originated predominantly from food samples and FCS, whereas isolates recovered during our sampling campaign were mainly derived from drains. Of these, 74 yielded complete allelic profiles, whereas two could not be assigned to a sequence type (ST) because some alleles remained unassignable due to poor sequence quality, despite repeated resequencing attempts and DNA re-extraction using the DNeasy Blood & Tissue Kit (Qiagen, Hilden, Germany).

Among the 74 typeable isolates, most belonged to lineage II (86.5%), while lineage I accounted for 13.5% (Fig. 8). Among them, 15 clonal complexes (CCs) and 21 STs were identified: the most frequent STs were ST9 (n = 23), ST37 (n = 6), and ST451 (n = 5). ST diversity differed between facilities, with F4 showing the highest diversity and harboring ten distinct sequence types. Several STs were repeatedly detected within the same facilities. Among the persistent STs (repeated isolation >6 months), ST204 was detected three times in F2, all from FCS. ST9 was detected 17 times in F3, predominantly from drains (12/17), but also from other non-food-contact surfaces, including the wheels of a smoking trolley (2/17) and the underside of a vat (1/17), as well as from FCS (2/17). In F4, ST87, ST121, and ST451, three, three, and five isolates were detected, respectively, and all corresponding isolates were recovered from FCS. ST9 was detected twice in F5, both times from drains. (Fig. 9). In contrast, repeated recovery of ST3406, ST236, ST517, ST37, ST9, and ST223 over shorter intervals (<6 months) was classified as recurrent detection rather than persistence.

**FIG 8.**
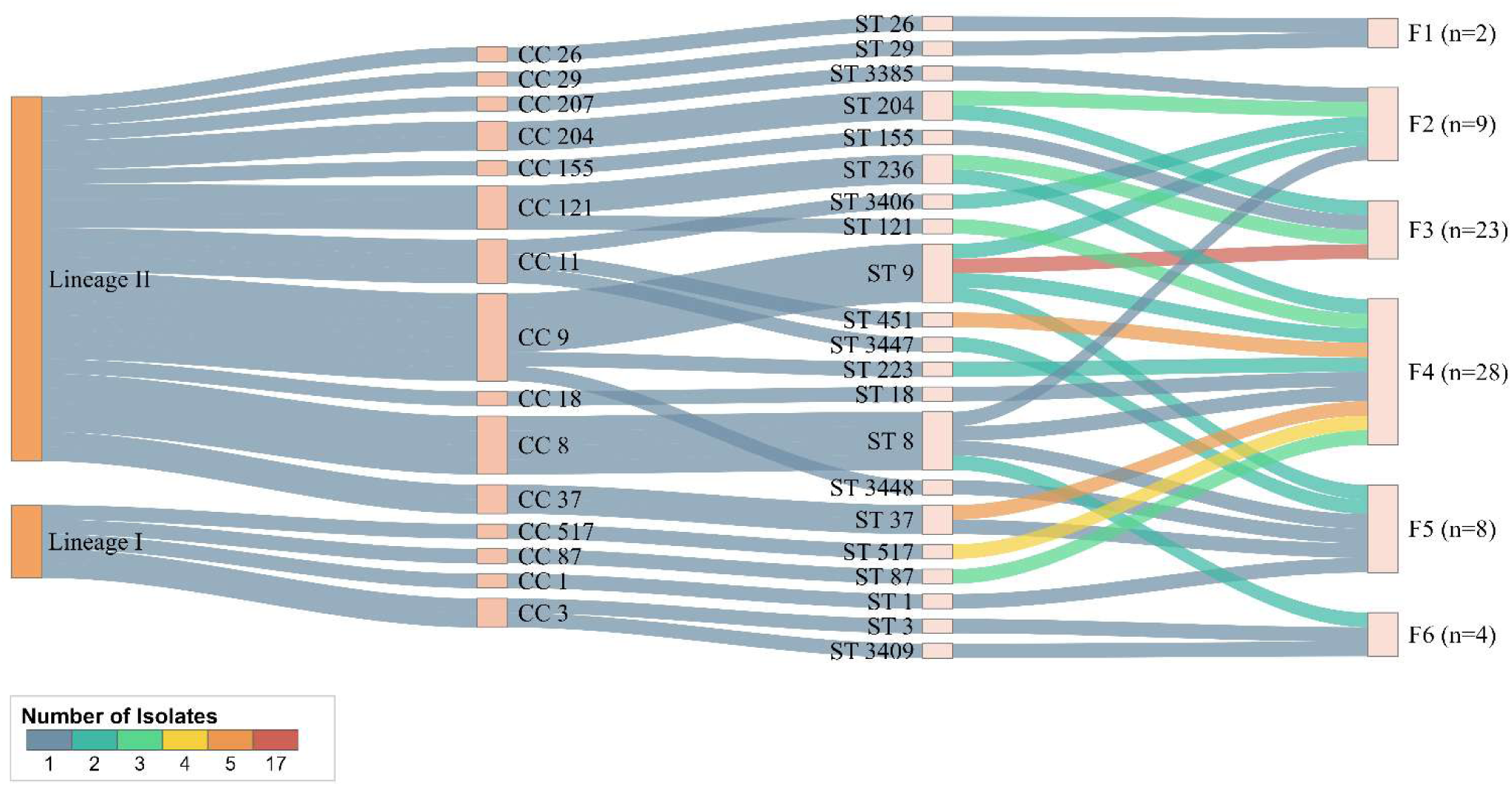
Distribution of *Listeria monocytogenes* isolates assigned by MLST across lineages, clonal complexes, sequence types, and facilities. Hierarchical alluvial diagram showing the distribution of the 74 typeable *L. monocytogenes* isolates across genetic lineages, clonal complexes (CC), sequence types (ST), and the six processing facilities. Colors represent how often a given sequence type was observed within a facility.

**FIG 9.**
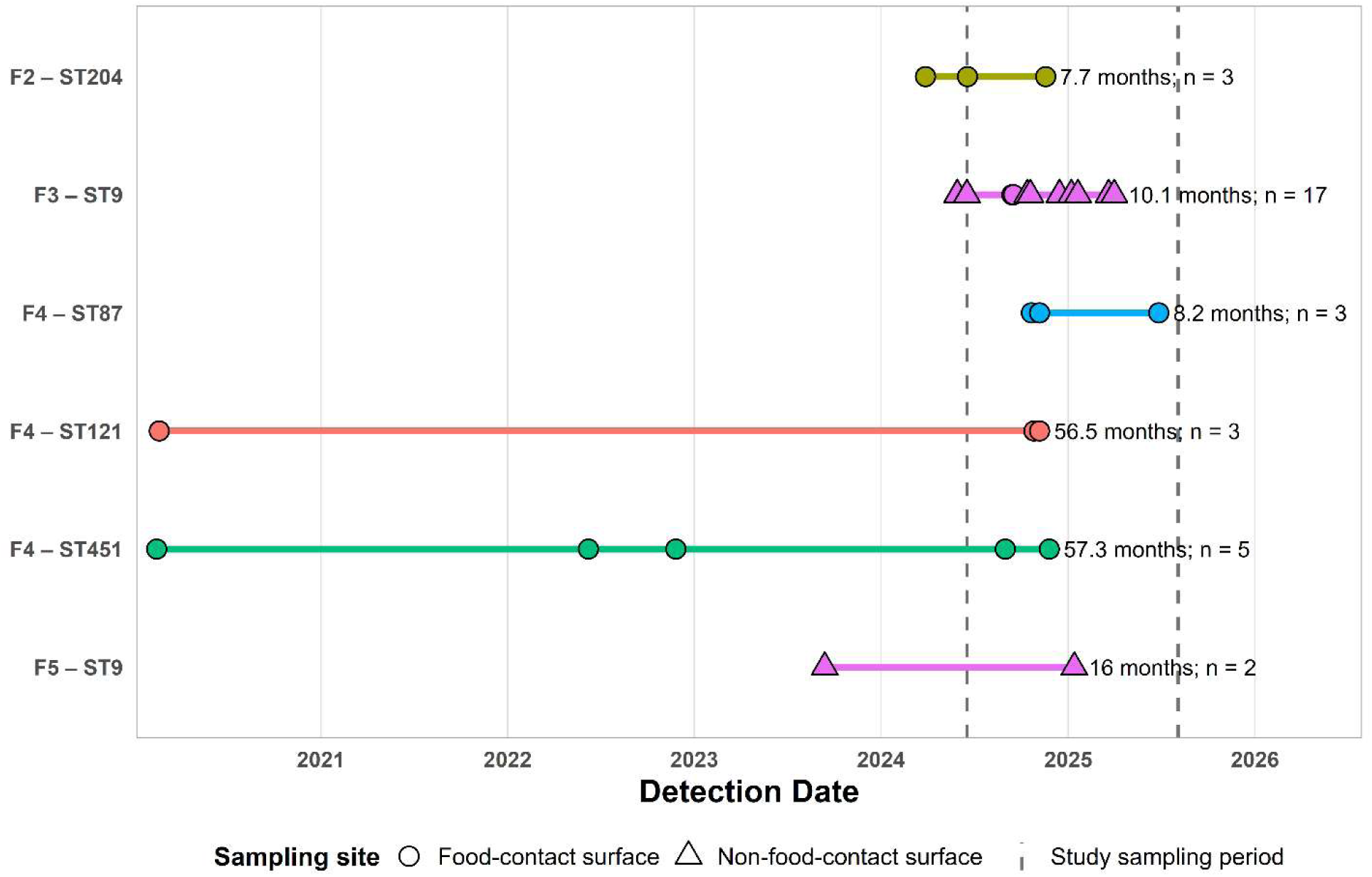
Detection time periods of persistent *Listeria monocytogenes* STs. Points indicate individual isolates, and horizontal lines indicate the period between first and last detection of each ST within the respective facility. Labels indicate the total detection period and number of isolates. Dashed vertical lines indicate the sampling period of the present study. Circles represent food-contact surface isolates, and triangles represent non-food-contact surface isolates.

## DISCUSSION

Across all facilities, genus-level abundance profiles were characterized by the repeated dominance of *Acinetobacter* spp. and *Pseudomonas* spp., which were the only genera identified as core genera across the entire dataset. Their broad distribution and high relative abundance suggest that these genera are particularly well adapted to environmental conditions present in meat processing facilities, including low temperatures, moisture, nutrient limitation, and repeated exposure to cleaning and disinfection. *Pseudomonas* spp. are frequently reported as dominant members of food-processing microbiota and include psychrotolerant, surface-associated, and biofilm-forming taxa that can persist under conditions typical of processing environments (Barcenilla et al., 2024; Bokulich et al., 2015; Tan et al., 2019). *Acinetobacter* spp. have also been repeatedly detected in food-processing environments and have been associated with cold, moist, and nutrient-limited niches, supporting their role as important members of resident FPE microbiota (Barcenilla et al., 2024; Xu et al., 2023). Together, the prevalence of these genera suggests that meat processing environments select for psychrotolerant and biofilm-associated microorganisms (Alves et al., 2024).

In addition to these shared core genera, several other taxa contributed prominently to facility-specific community profiles, including *Rhodococcus*, *Psychrobacter*, *Chryseobacterium*, *Flavobacterium*, and *Aeromonas* species. Their distribution varied between facilities indicating that local conditions such as raw material input, moisture availability, surface properties, temperature, and sanitation efficiency shape the composition of resident microbial communities. Many of these genera have also been reported from cold, moist, or nutrient-limited food-processing environments and include taxa capable of surface association and biofilm formation (Møretrø and Langsrud, 2017; Wagner et al., 2021), for example *Flavobacterium* spp. on solid surfaces (Magesh et al., 2024) or *Aeromonas* spp. on glass and polystyrene (Ormanci and Yucel, 2017). Thus, while *Acinetobacter* spp. and *Pseudomonas* spp. represented broadly distributed core members, the additional predominant genera likely reflect facility-specific ecological selection within individual processing environments.

Despite shared dominant taxa, community composition differed between facilities, indicating distinct resident microbial communities in individual facilities. This variation likely reflects differences in facility characteristics, such as size, hygiene practices, and processed animal species. Within-facility variation further indicates that the local microenvironment, influenced by surface properties, moisture, and cleaning efficiency, shapes microbial communities (Belk et al., 2022; Xu et al., 2025).

Core microbiota analysis showed that microbial community stability depended on taxonomic resolution and facility context. At the genus level, *Pseudomonas* spp. and *Acinetobacter* spp. formed a shared core across all facilities, indicating common environmental selection pressures in meat processing environments. In contrast, no ASV was shared as a core taxon across all facilities, and no core ASVs were identified in F2 or F3, although both facilities contained core genera (*Pseudomonas* and *Acinetobacter* species). This suggests that similar ecological niches may be occupied by different ASVs across facilities or over time, resulting in genus-level stability but ASV-level turnover. This may also be reflective of the similar growth requirements between ASVs of the same genus. Similar genus-level stability together with finer-scale taxonomic variability has been reported in other food-processing environments (Asmus et al., 2025; Barcenilla et al., 2024). Facilities lacking core ASVs, such as F2 and F3, may therefore represent more temporally dynamic microbial communities. This interpretation is consistent with the temporal analyses, which showed significant changes in Shannon diversity in F2 and F3 and significant longitudinal community change in F2. Such community turnover may reduce the likelihood that individual ASVs persistently meet core-microbiota criteria across sampling events. The temporal analyses further indicated that changes in Shannon diversity and longitudinal community composition did not follow a simple monotonic trend over the study period. Instead, the better fit of models treating sampling event as a categorical predictor suggests event-specific, non-linear temporal dynamics. These patterns may reflect episodic or facility-specific influences, such as variation in raw material inputs, moisture levels, sanitation efficacy, or microbial recolonization after cleaning, rather than a uniform directional trend over time.

These patterns may also provide ecological context for *Listeria* spp. occurrence. Facilities F2 and F3 had the highest numbers of *L. monocytogenes* detections in the present study, corresponding to detection prevalences of 8.6% in F2 (6/70 samples) and 10.0% in F3 (6/60 samples). Because most *Listeria* spp.-positive samples were recovered from drains, this pattern should be interpreted in the context of wet, difficult-to-sanitize niches. Temporally dynamic communities in wet or difficult-to-sanitize areas may indicate repeated microbial input, disturbance after sanitation, or changing nutrient and moisture conditions, all of which could create opportunities for

*Listeria* spp. introduction, survival, or repeated recovery. Conversely, more stable communities, such as those observed in cold rooms, may reflect more constant environmental conditions, lower rates of microbial introduction and turnover, or less frequent disturbance. However, whether community stability inhibits or favors *Listeria* spp. cannot be inferred from community data alone, because stable communities may either resist invasion or provide persistent niches that support *Listeria* spp. survival.

Notably, production areas were distinguished more strongly by community composition than by alpha diversity, suggesting that functional zones differed primarily in the identity of their resident taxa rather than in overall within-sample diversity. This likely reflects area-specific selection pressures, including differences in moisture, temperature, organic residue availability, surface properties, and cleaning efficiency, which shape the composition and temporal stability of microbial communities in different production areas (Belk et al., 2022; Botta et al., 2020).

Together, these findings indicate that *Listeria* spp. occurrence should be interpreted in the context of facility-specific and temporally dynamic microbial communities rather than as the result of single taxa alone.

Culture-based analyses revealed a pronounced site-specific distribution of *Listeria* spp. The higher detection frequency in drains than on FCS highlights the importance of wet, difficult-to-sanitize niches, which can promote biofilm formation and persistence of *Listeria* spp. (Fagerlund et al., 2017). Similar site-dependent patterns have been described in other FPEs (Barcenilla et al., 2024). This observation is consistent with the view that hygienically challenging environmental niches contribute to *Listeria* spp. presence in FPEs (Hazards (BIOHAZ) et al., 2024). By including both FCS and drains, the study provides a differentiated view of contamination patterns within FPEs.

ASV-level associations in drain samples should be interpreted as ecological correlations rather than evidence of direct interactions. The positive associations of ASVs assigned to *Acinetobacter*, *Rhizorhapis*, and *Vagococcus* species with *Listeria* spp.-positive drain samples may reflect shared preferences for wet, nutrient-rich, or biofilm-associated niches that also support *Listeria* spp. survival or repeated recovery. Genera such as *Acinetobacter* spp. (Burnett et al., 2025; Pracser et al., 2024) and *Flavobacterium* spp. (Belk et al., 2022) have previously been reported in *Listeria* spp.-positive environments. It is also possible that *Listeria* spp. profit from the protected microenvironment that biofilms built by prolific biofilm producers like *Acinetobacter* spp. provide (Møretrø and Langsrud, 2017). Conversely, ASVs showing negative, non-significant trends may reflect habitat separation or environmental conditions less suitable for *Listeria* spp., such as drier surfaces, lower nutrient availability, or more effective sanitation, rather than direct inhibition. At the same time, a direct antagonistic effect is also plausible for some negatively associated taxa, for example, through bacteriocin-producing bacteria (LAB) such as *Lactococcus* spp., *Streptococcus* spp., *Lactobacillus* spp., and *Enterococcus* spp., which can suppress *L. monocytogenes* (Camargo et al., 2018). Experimental co-culture and multi-species biofilm studies could determine whether these taxa directly affect *Listeria* spp. fitness, survival, or sanitizer tolerance.

*Listeria* spp. detection was performed according to ISO 11290, ensuring comparability with routine monitoring data. Nevertheless, enrichment-based detection is not equally sensitive for all *Listeria* spp. and may underestimate overall species diversity, particularly for less frequently recovered taxa such as *L. seeligeri*, *L. grayi*, and potentially *L. ivanovii* (Carlin et al., 2022). *L. seeligeri* and *L. grayi* were detected in our study despite their potentially reduced recoverability. However, it is plausible that additional occurrences of these species were missed in other samples. Accordingly, our culture-based data likely provide a robust estimate of *Listeria* spp. occurrence but may not fully reflect the true species composition or relative abundance of *Listeria* spp. in sampled facilities.

MLST analysis revealed identical *L. monocytogenes* STs at product-associated sampling points, FCS, and floor drains within the same facilities. Recovery of isolates with identical STs from these sites is consistent with persistence, repeated introduction, or transfer within the same processing environment (Fagerlund et al., 2022). Drain contamination may therefore provide useful information on local conditions and on the occurrence of *L. monocytogenes* lineages repeatedly recovered within a processing room (Belias et al., 2022; Dzieciol et al., 2016).

The predominance of lineage II may be explained by lineage-specific ecological differences. Lineage I is often associated with clinical cases and higher virulence potential, whereas lineage II is more frequently recovered from food and food-processing environments and includes clonal complexes repeatedly linked to environmental persistence (Orsi et al., 2011). Accordingly, lineage II strains may be favored in FPEs because they are well adapted to refrigeration temperatures, nutrient limitation, surface-associated growth, and repeated cleaning and disinfection. This interpretation is consistent with the frequent detection of CC9, CC121, and CC204 in the present study, which have been repeatedly associated with meat processing environments and persistence (Martín et al., 2014; Voronina et al., 2023).

In our study, repeated recovery of specific STs within facilities met our definition of persistence. These STs belonged to CC9, CC11, CC121, and CC204, clonal complexes that include genotypes frequently reported from FPEs. While MLST does not provide sufficient resolution to infer persistence of identical strains, the repeated detection of STs within these clonal complexes supports the interpretation that these specific STs are well adapted to meat processing environments. This is consistent with previous studies reporting persistent or recurrent *L. monocytogenes* in food-processing environments, particularly among isolates belonging to CC9, CC121, and CC204 (Fagerlund et al., 2022; Guidi et al., 2021), as well as studies from meat processing facilities where these clonal complexes have repeatedly been associated with persistence (Martín et al., 2014; Stoller et al., 2019). In addition, *L. monocytogenes* strains belonging to these recurrent clonal complexes have been associated with increased stress tolerance, survival after cleaning and disinfection, and the ability to colonize environmental niches and form biofilms (Fagerlund et al., 2022; Stoller et al., 2019).

Altogether, this study provides a longitudinal view of microbial community structure and *Listeria* spp. occurrence in six German meat processing facilities by integrating microbial community profiling, culture-based detection, and MLST over one year. Our findings show that *Listeria* spp. recovery should be interpreted within the ecological context of facility-specific resident microbiota, wet and difficult-to-sanitize niches such as floor drains, temporal community dynamics, and co-occurring taxa associated with *Listeria* spp.-positive samples. This perspective moves environmental *Listeria* spp. surveillance beyond presence–absence detection, highlighting the importance of considering the microbial community background and niche characteristics of sampling sites. While routine monitoring should continue to prioritize high-risk sites such as FCS, microbiome-informed approaches may provide complementary information by helping to identify environmental niches with microbial community characteristics associated with *L. monocytogenes* survival or repeated recovery. Repeated recovery of identical *L. monocytogenes* STs further supports the relevance of facility-associated reservoirs. In line with previous work showing that food-processing environments harbor facility-associated microbiomes linked to *Listeria* spp. occurrence (Belk et al., 2022) and diverse, facility-specific meat processing microbiomes (Barcenilla et al., 2024), our results emphasize that *Listeria* spp. surveillance should be interpreted within the broader microbial ecology of each facility.

## DATA AVAILABILITY

Sequencing data are available in the NCBI under BioProject PRJNA1261676. Metadata and Sequencing data analysis code is available at: https://github.com/kovaclab/environmental_microbiota_meat_processing_facilities.

## ACKNOWLEDGMENTS

We thank the Institut Pasteur teams for the curation and maintenance of BIGSdb-Pasteur databases at https://bigsdb.pasteur.fr/. We thank Julia Mühmel and Meike Schumann for preparing the selective media used for *Listeria* spp. enrichment and isolation. We further thank Barbara Fritz, Erika Altgenug, and René Mamet for their assistance with MALDI-TOF identification of isolates. During the preparation of this manuscript, the authors used ChatGPT-5.5 to improve readability and language. After using this tool, the authors reviewed and edited the content as needed and took full responsibility for the final publication. JK was supported by the USDA National Institute of Food and Agriculture and Hatch Appropriations under Project PEN04853 and Accession 7005519, and the Multistate project 4666, as well as the Lester Earl and Veronica Casida Career Development Endowment. JK’s work at University of Munich was supported by the DAAD scholarship and Research Fellowships of the Alexander von Humboldt Foundation.

